# D-amphetamine alters the dynamic ECoG activity distribution patterns in the rat neocortex

**DOI:** 10.1101/2025.05.09.653067

**Authors:** Astrid Mellbin, Henrik Jörntell, Fredrik Bengtsson

## Abstract

Amphetamine has widespread effects on multiple neurotransmitter systems, potentially altering the physiological connectivity and network dynamics across various regions of the brain. In this study, we investigated the effects of D-amphetamine using our previously published approach where electrocorticogram (ECoG) recordings from eight cortical areas provided a coarse estimation of the global activity distribution patterns across sets of neuron populations. Changes in these activity distribution patterns were quantified with Principal Component Analysis (PCA) and k-Nearest Neighbors (kNN) classification. We found that D-amphetamine significantly altered the activity distribution patterns both for spontaneous activity and for activity recorded during ongoing tactile stimulation. It also reduced the difference between spontaneous activity and activity during ongoing tactile stimulation, which suggests that amphetamine reduced the organization in the network activity and could potentially explain hallucinations under the influence of amphetamine. Each of these changes were distributed approximately evenly across each dimension of the principal component space. This indicates that amphetamine impacts cortical network dynamics broadly and in multifaceted ways, compatible with the system-wide presence of the receptors that amphetamine interferes with. Our data indicates that relatively low doses of D-amphetamine can induce changes in brain activity distributions that are measurable potentially also by non-invasive EEG electrodes.

## Introduction

The idea of a globally interconnected functional network in the neocortex has in recent years gained increased recognition. Studies have shown that tactile information is decoded in several different cortical areas and that information from visual input in awake mice can be found across the cortex[1–3]. Recent wide-field calcium imaging studies also indicate that there is a global activation of the cortex across a variety of behavioral actions[4]. Motivated by a need for more non-invasive methods to explore global activity distributions, we recently showed that a method designed to analyze global changes in cortical activity distributions can be used to detect even weak tactile inputs using the less invasive recording techniques of electroencephalogram (EEG) or electrocorticogram (ECoG)[5].

If the cortex operates as a globally interconnected network, any damage or disruption to the cortex would be expected to alter the activity and the processing of the whole network. For example, a stroke in a remote cortical area decreases the ability of the neurons in primary somatosensory cortex (S1) to process tactile information[6]. It is reasonable to assume that if a small, localized lesion can impact cortical processing at such a distance, then more widespread disruptions could potentially have bigger effects on the cortical network and its processing capabilities. Amphetamine is a drug known to have a widespread effect on the brain, affecting the neurotransmitters noradrenaline, dopamine, acetylcholine and serotonin which through axonal projections from the brainstem impact essentially all parts of the cortex and thalamus[7,8]. Amphetamine can be used to treat ADHD and narcolepsy[9] and has therefore been examined for some of its effects on the cortex. D-amphetamine has been found to impact the frequency content of EEG in a way that suggested an activation of D1-receptors, with a switch to activation of D2-receptors with repeated administration[10]. Another study found evidence of amphetamine causing forebrain arousal by acting on noradrenergic β-receptors[11]. Amphetamine also increases the release of acetylcholine in the cortex, by a mechanism that appears to depend on more than just an activation of D1- and D2-receptors[12]. When amphetamine is misused as a drug, users can exhibit symptoms from a wide range of modalities, such as extreme moods, ataxia, stereotypical mouth movements, increased sympathetic stimulation and paranoia[13].

Other studies focused on the effect of amphetamine on the functional connectivity using fMRI, a method that reports the spatial features of the brain activity integrated over time. A reduction in functional connectivity was found in the cortico-striato-thalamic network, as well as in the default mode networks and the salience-executive networks[14]. Other studies have reported a reduced functional connectivity between nucleus accumbens and the basal ganglia, medial prefrontal cortex, temporal cortex, and the anterior cingulate cortex, with an increase in functional connectivity between nucleus accumbens and medial frontal regions as well as between putamen and the left inferior frontal gyrus[15,16]. Furthermore, D-amphetamine induces an auditory-sensorimotor-thalamic functional hyperconnectivity measured with fMRI[17]. In the cortex, amphetamine has been shown to reduce both REM and non-REM sleep times in rodents, while also reducing low frequency EEG activity[18]. Amphetamine was also reported to modulate synaptic plasticity in the motor cortex, allowing better task specific recovery after brain lesions, and speeding up the learning of motor tasks[19,20]. However, another study found that while amphetamine increased short lasting neuronal excitability, it suppressed long lasting plasticity induced by stimulation[21].

However, as the dynamic global collaboration between the neurons appears to be a critical aspect of brain operation, it follows that methods designed to quantify the dynamically changing distributions of global activity may potentially provide for sensitive indicators also of more subtle, but systematic, changes in brain activity. Since a high temporal resolution is a key to address activity dynamics, electrophysiological recording methods remain advantageous in this regard[22]. But given that it is impossible to record the electrical activity of every single neuron in the brain, mass recordings such as electrocorticogram (ECoG) across multiple electrodes may be useful to provide insights into systematic shifts in the dynamic activity distribution across the global network[5] and provide results that are potentially translatable to humans. Here we used the same methodology to examine how D-amphetamine affects the global ECoG activity distribution patterns[5]. Using PCA and kNN analysis, we find that D-amphetamine significantly alters the activity distribution patterns both for spontaneous activity and for activity recorded during ongoing tactile inputs. We also find that it reduces the difference between spontaneous activity patterns and activity patterns during ongoing sensory inputs, suggesting a general disorganization of the dynamic structure of the brain network activity.

## Results

### D-amphetamine induces changes in the brain activity distribution patterns

We used a set of eight ECoG electrodes distributed globally across the cortex as shown in Figure 1A. The ECoG activity was continually recorded during the stimulation protocol (Figure 1B-D), which contained periods of spontaneous activity alternated with periods with ongoing tactile stimulation to the second digit of the forepaw or the hind paw (Figure 1B). To investigate the impact of amphetamine on the brain activity distribution patterns, we compared the brain activity data both for the spontaneous activity and for the activity with ongoing stimulation (Figure 2A). Using principal component analysis (PCA) of the ECoG activity distribution patterns across the eight ECoG electrodes, each time step of the recorded activity was mapped to the PC-space (Figure 2B), in which the clustering of the data for the two different conditions in the comparison was quantified using the k-Nearest Neighbor (kNN) method. Across the 7 experiments, each with 7 stimulation episodes at different frequencies for two different stimulation sites, a total of 98 different comparisons were made (both for spontaneous activity and for activity during ongoing tactile stimulation). We first quantified the changes in the spontaneous activity induced by D-amphetamine (Figure 2C). As a control, we also compared different half-segments of spontaneous activity under each condition (with and without D-amphetamine, respectively) with each other (Figure 2A,C). The kNN accuracies of the difference in activity induced by D-amphetamine were significantly higher than those obtained for the control data (Friedmans <0.01, post hoc sign test <0.01 against both sets of control data). Similar results were obtained for the changes D-amphetamine induced in the activity with ongoing tactile stimulation (Figure 2D, data from forepaw and hind paw stimulations combined) (Friedmans <0.01, post hoc sign test <0.01 against both sets of control data). The results are summarized in Table 1.

**Figure 1.**
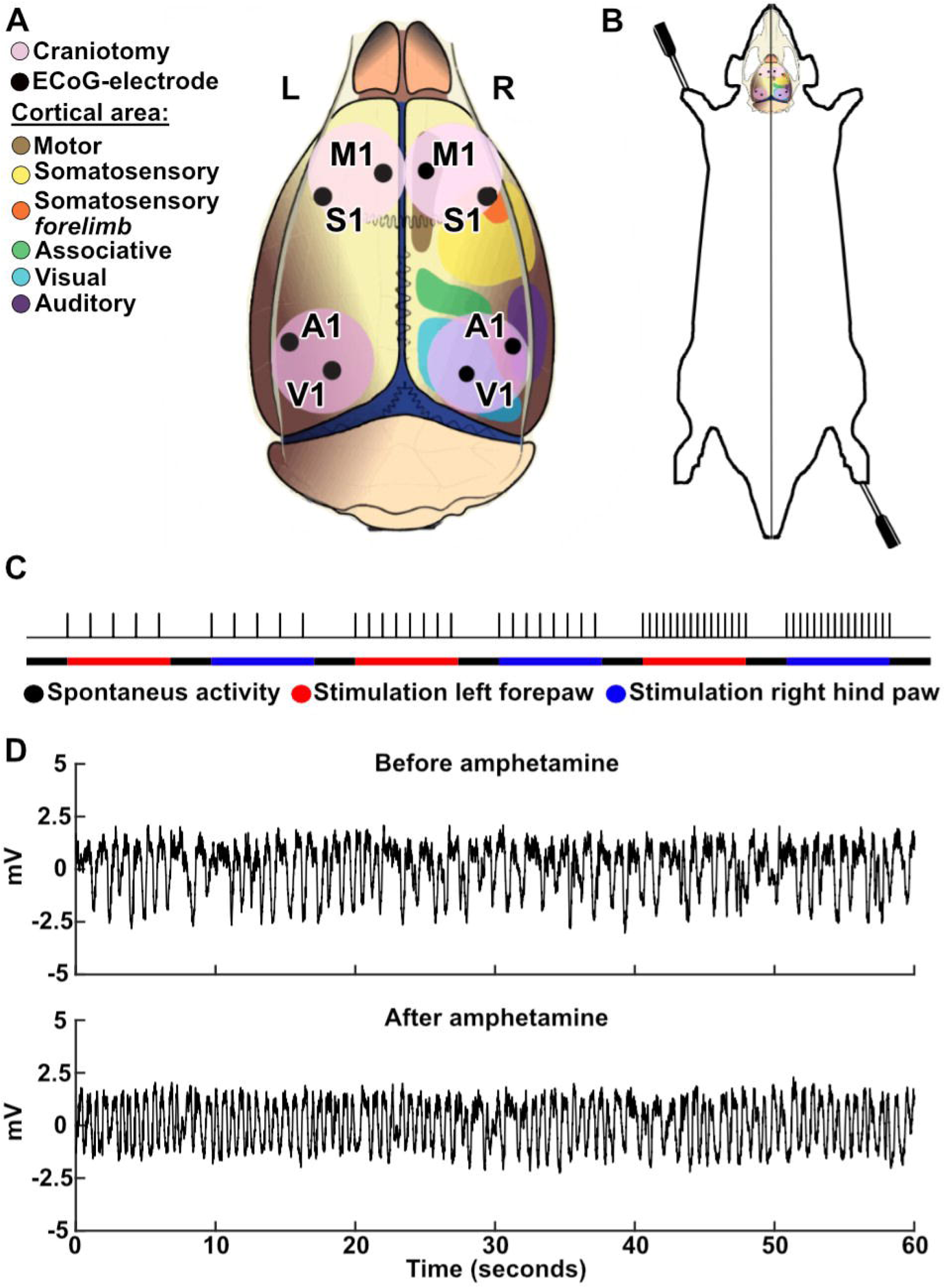
Recording setup and stimulation protocol. **A** The location of craniotomies and recording electrodes in relation to different cortical areas. M1, primary motor cortex; S1, primary somatosensory cortex; A1, primary auditory cortex; V1, primary visual cortex. **B** The location of the tactile stimulation electrodes on the distal left forepaw and right hind paw. **C** Visualization of the stimulation protocol that was repeated before and after amphetamine administration. **D** ECoG traces before and after amphetamine administration, recorded from the right S1 area, with artifacts removed and Savitzky-Golay filter applied. The traces were recorded one minute before (top) and 15 minutes after the administration of amphetamine (bottom).

**Figure 2.**
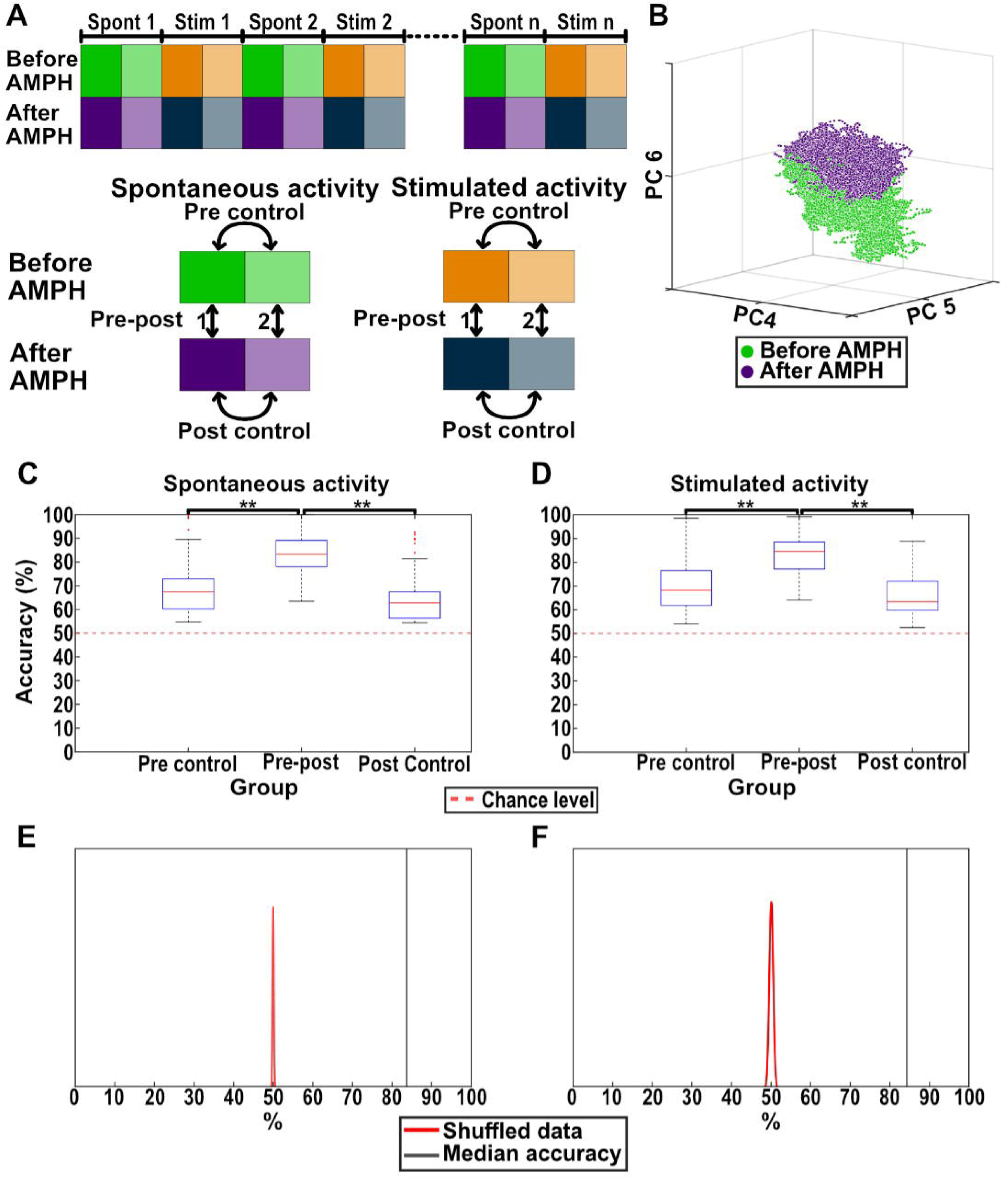
Both spontaneous and stimulated activities were altered by D-amphetamine. **A** Illustration of the comparisons made in the kNN analysis. To make comparisons between matching periods of the protocol, each period of spontaneous and stimulated activity was divided into two halves (dark and light nuances of the same color). The two halves were compared against each other, both for the spontaneous and the evoked activity, to obtain control values (pre-control’ and ‘post-control’). These control values were then compared to the activity differences obtained after amphetamine (‘pre-post 1’ & ‘pre-post 2’, which were combined into one ‘Pre-post’ value, to ensure similar size of the data sets used in kNNs that are directly compared). **B** Distribution of the spontaneous activity in principal component space (the subspace defined by PCs #4-6) before and after the administration of D-amphetamine in a sample experiment. **C** The kNN accuracy of the comparison between the spontaneous activities before and after D-amphetamine administration (‘Pre-post’). Also shown are the kNN accuracies from the control comparisons before and after D-amphetamine administration (‘Pre-control’ and ‘Post-control’). Asterisks indicate significantly different distributions at p<0.01 (Sign test). Each box with outliers shows all 98 kNNs for the group. **D** Similar to **C** but for activity recorded during stimulation periods. Data from forepaw and hind paw stimulations are combined. **E** Results from the data shuffling. Illustration shows the results from one of the 98 kNNs, chosen for being the comparison with an accuracy closest to the median. The red distribution curve represents the results for the shuffled data (the shuffling was repeated 50 times), and the black line shows the kNN accuracy in the test data (pre-post’). **F** Similar to **E** but for stimulated activity.

**Table 1.** kNN accuracies for the differences induced by amphetamine in the spontaneous and in the stimulated activities. The first column shows the test result, i.e. the median kNN accuracies obtained for the differences between the activity data before and after D-amphetamine. The periods of spontaneous activity and the periods of stimulated activity are investigated separately. The second column shows the difference between the two halves of data recorded before D amphetamine (‘Pre control’) and the third column shows the corresponding control data for the two halves of data recorded after D-amphetamine (‘Post-control’).

|  | Difference between before and after D-amphetamine (Pre-post) (%) | Control before D-amphetamine (Pre control) (%) | Control after D-amphetamine (Post control) (%) | N (stimulation periods) |
| --- | --- | --- | --- | --- |
| **All, spontaneous activity** | 83,14 | 67,36 | 62,72 | 98 |
| **All, stimulated activity** | 84,55 | 68,16 | 63,34 | 98 |

As an additional control, we used data from 5 experiments, which had the same total recording durations, but without the D-amphetamine administration. The kNN accuracy of the difference between two consecutive time segments (two hours) of spontaneous activity were significantly worse than for the two segments of spontaneous data with and without D-amphetamine (Wilcoxon rank sum test p<0.01), indicating that D-amphetamine significantly altered the spontaneous activity patterns also relative to this control.

Using data shuffling, we found that the actual chance level for the kNN analysis was centered around the theoretical chance level of 50%, i.e. that the kNN could not detect any difference between the normal activity and the activity under the D-amphetamine regime when the conditions (labels) of the data points were shuffled (Figure 2E-F). Moreover, the shuffled data had a very narrow distribution and the actual data was located many standard deviations away from the shuffled data (Figure 2E-F).

### D-amphetamine reduced the difference between spontaneous and stimulated activity

As we have previously reported, the analysis of the global ECoG activity distributions can be used to detect differences between spontaneous activity and the activity when there is a weak ongoing tactile stimulation (Mellbin et al, 2024). We repeated the same analysis here but in addition compared the difference between spontaneous and stimulated activity after D-amphetamine administration (Figure 3A). Under both conditions (with and without D-amphetamine) the kNN accuracy of the actual data was substantially different from distribution of the shuffled data (Figure 3B,C). Across the whole data set, including when the forelimb and the hindlimb stimulations were considered separately, the results were equivalent (Figure 3D, left) also under D-amphetamine (Figure 3D, right). However, when compared against the shuffled data, the difference between spontaneous and stimulated activity was smaller after D-amphetamine (Figure 3B, C). Indeed, the kNN accuracy was significantly higher before the administration of D-amphetamine compared to after, for both forepaw and hind paw stimulation (p <0.01 for either stimulation) (Table 2). This indicates that D-amphetamine reduced the difference between spontaneous and stimulated activity. We also quantified whether the difference between the spontaneous and stimulated activity was significantly different depending on whether forepaw or hind paw stimulation was used. We found that it was not, neither before (p =0.25) nor after (p =1) D-amphetamine.

**Figure 3.**
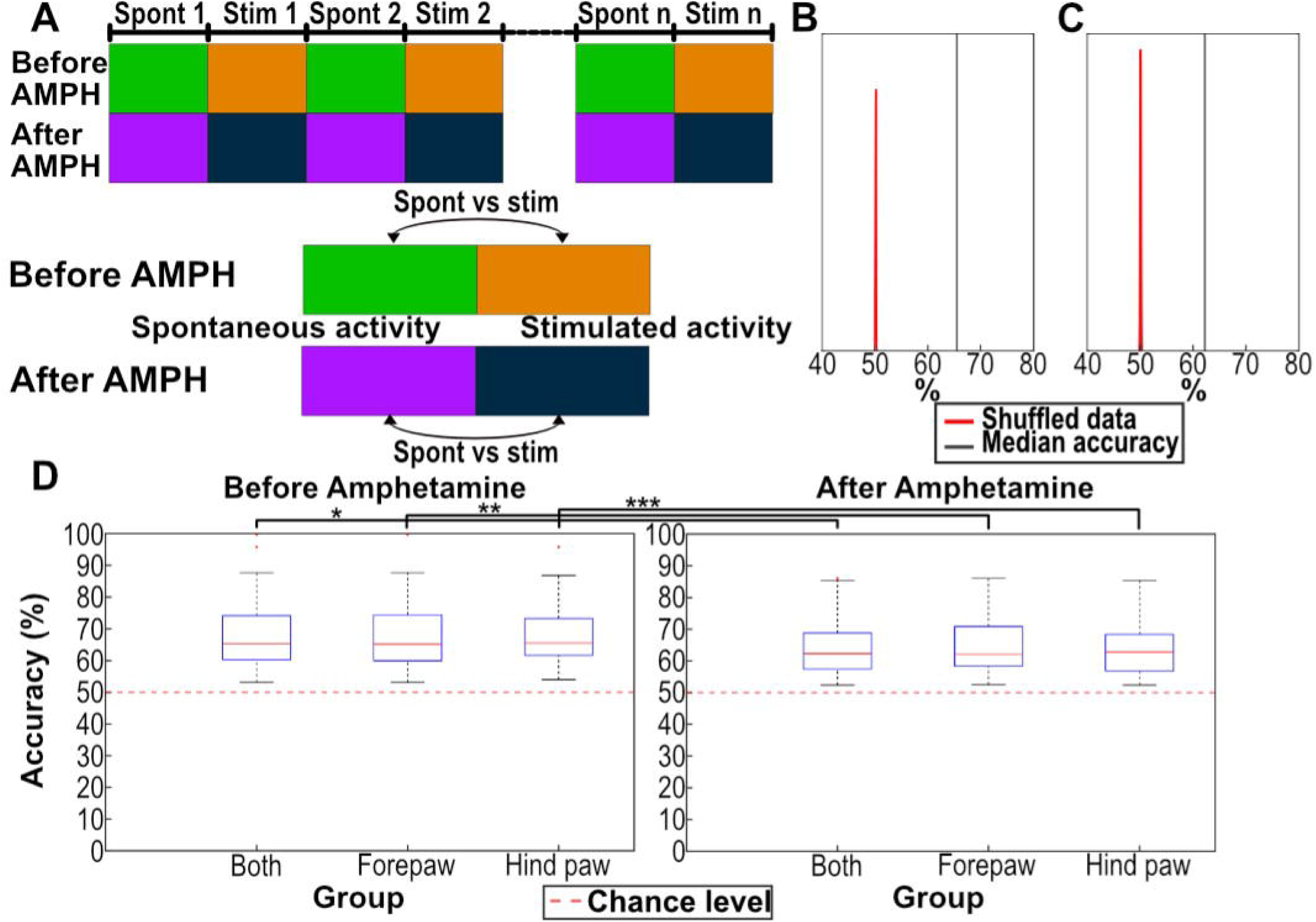
The difference between spontaneous and stimulated activity was reduced by D-amphetamine. **A** Illustration of the comparisons made in the kNN analysis. Spontaneous and stimulated activity was compared to each other, before and after administration of D-amphetamine, respectively. **B** kNN results compared to the shuffled data. The result (black line) represents the median accuracy for one of the 98 kNN analyses made, chosen for being the comparison with an accuracy closest to the median accuracy of all the 98 kNN results. The red curve represents the kNN results for the shuffled data, the shuffling being repeated 100 times. **C** Similar to **B** but for data after amphetamine administration. **D** The kNN accuracy of the difference between spontaneous and stimulated activity for the activity recorded before and after D-amphetamine administration (left and right diagram, respectively. Comparisons were made both for activity recorded during stimulation of the forelimb and for the activity recorded during stimulation of the hindlimb, as well as the activity recorded under either stimulation (‘Both’). In “Both” the box with outliers represent the result of all 98 kNN analyses, whereas “Forepaw” and “Hind paw” data included 49 kNN analyses each.

**Table 2.** kNN accuracies for the difference between stimulated and spontaneous activity induced by D-amphetamine.

|  | Spontaneous<br>vs Stimulated,<br>before<br>amphetamine<br>(%) | Spontaneous<br>vs Stimulated,<br>after<br>amphetamine<br>(%) | N<br>(stimulation<br>periods) |
| --- | --- | --- | --- |
| **All** | 65,39 | 62,33 | 98 |
| **Stimulation**<br>**location, paw** |  |  |  |
| **Forepaw left** | 65,17 | 62,13 | 49 |
| **Hind paw**<br>**right** | 65,53 | 62,85 | 49 |

The columns show the category of the comparison and the rows show the results based on which stimulations were included in the kNN analysis (the results represent the median accuracy across all experiments).

### The dimensionality of the brain activity was not affected by D-amphetamine

Our basic analysis approach was to use PCA to map the brain activity distribution recorded at each time step to a location in the high-dimensional space defined by the PCs and then to calculate the Euclidean distances between the data points to obtain a kNN value. To explore if D-amphetamine affected the dimensionality of the data, the kNN accuracy was iteratively calculated based on subsets of the PCs. We first examined the contribution of each PC to explain the alteration induced by D-amphetamine in the spontaneous activity and in the stimulated activity (Figure 4A,B). The accuracy for separating the control condition from the D-amphetamine condition increased for each added PC, for both spontaneous and stimulated activity (Figure 4A). Likewise, removing any PC decreased the kNN accuracy for both types of activity (Figure 4B). Hence, in this case the effect of D-amphetamine appeared to be broadly distributed across all dimensions of the brain activity data. This indicates that D-amphetamine did not alter the dimensionality of the brain activity distribution patterns or bias the activity to any specific such dimension.

**Figure 4.**
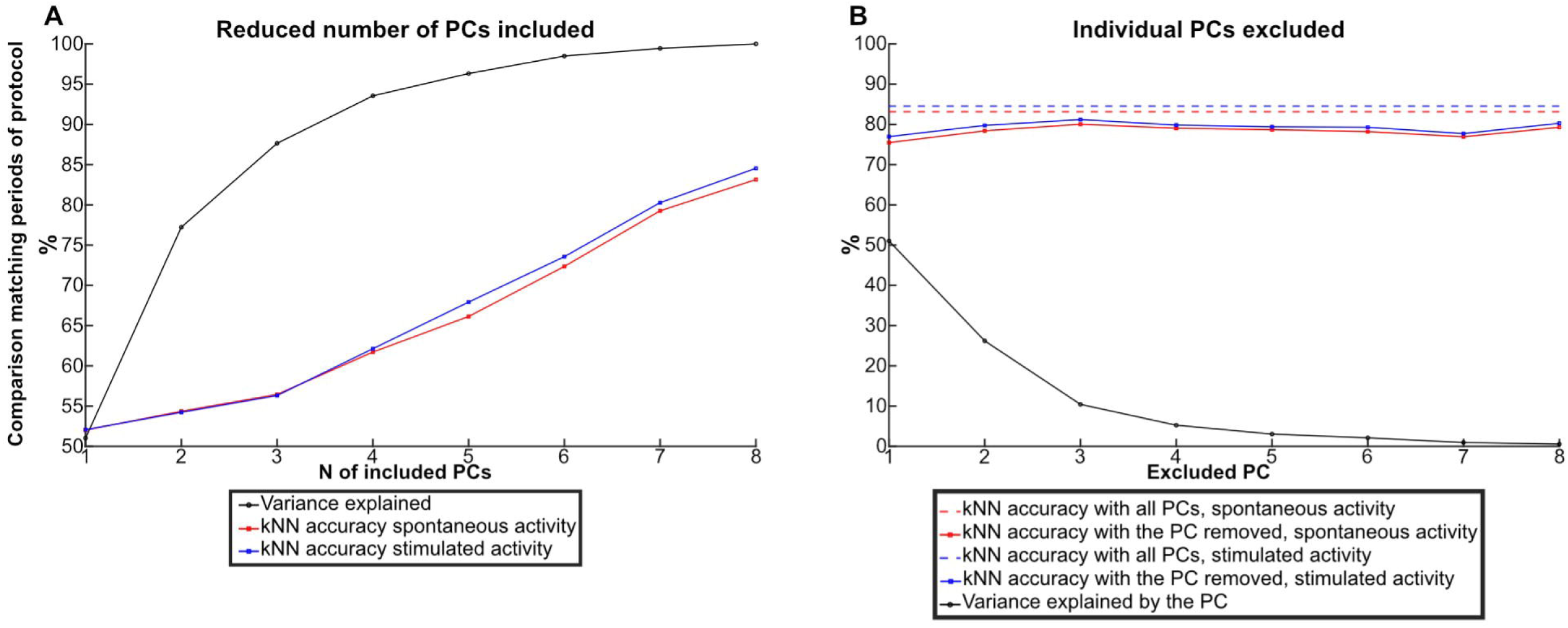
Each PC contributed to the measured difference induced by D-amphetamine in both the spontaneous and the stimulated activity. **A** The median kNN accuracy (from 98 kNNs) of the difference in the activity data induced by D-amphetamine as a function of the number of included PCs in the kNN analysis. Red data points show the median kNN accuracy for the data recorded during spontaneous activity (before and after D-amphetamine) and blue data points show the median kNN accuracy for data recorded during stimulated activity (before and after D-amphetamine). The median cumulative variance (from seven rats) explained as a function of the number of PCs is shown by the black data points. **B** Dashed lines indicate the median kNN accuracy (from 98s kNNs) for separating the activity data with and without D-amphetamine for both spontaneous and stimulated activity. Solid lines and data points indicate the medina kNN accuracy per each PC removed from the analysis. Black data points indicate the median variance explained (from seven rats) by each PC.

We next analyzed how the difference between spontaneous and stimulated activity was altered by D-amphetamine. Figure 5A shows that each PC added to the kNN accuracy also after D-amphetamine, again suggesting that D-amphetamine did not impact any specific aspect of the brain activity distribution patterns. However, the effect of each added PC was reduced after the administration of D-amphetamine (sign test p <0.01, all 98 kNN accuracies included). This suggests that the structure of the ECoG activity data was impacted by the drug, making it harder for the kNN analysis to separate the spontaneous and stimulated activity after D-amphetamine. Figure 5B instead analyzed if any specific PC had a particularly large importance for explaining the difference between the spontaneous and stimulated activity. But like Figure 4B, we found that any PC removed affected the resulting accuracy, both before and after D-amphetamine administration. Moreover, in neither case did the magnitude of the reduction in accuracy correlate with how much of the variance the PC explained (Pearson’s correlation coefficient -0.22 pre administration, -0.06 post administration).

**Figure 5.**
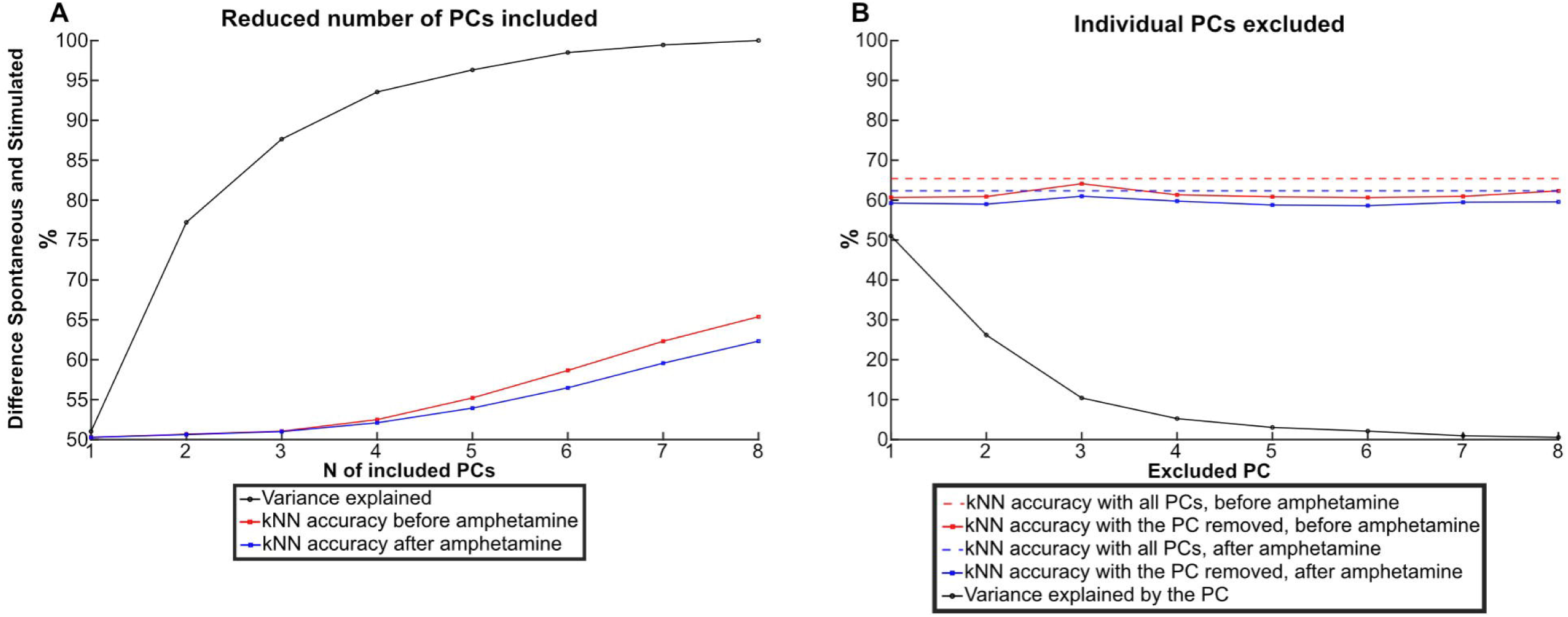
Each PC contributed to the difference between spontaneous and stimulated activity, both before and after D-amphetamine. **A.** The median kNN accuracy (from 98 kNNs) of the difference between the spontaneous and the stimulated activity data as a function of the number of included PCs. Red data points show the median kNN accuracy for data recorded before D-amphetamine and blue data points show the median kNN accuracy for data recorded after. The median cumulative variance explained (from seven rats) per PC for the entire data set is shown by the black data points. **B.** Dashed lines indicate the median kNN accuracy (from 98 kNNs) of the difference between the spontaneous and the stimulated activity data. Solid lines and data points indicate the median kNN accuracy per each PC removed from the kNN analysis. Black data points indicate the median variance explained (from seven rats) by each PC.

### D-amphetamine caused a general decrease in power across frequency bands

We also analyzed the power across the frequency bands of the ECoG signal before and after the administration of D-amphetamine. A significant decrease in power was found for all frequency bands after the D-amphetamine administration (Sign test, p<0,01). This was true also when the data was split into spontaneous and stimulated activity, with the exception of the Beta frequency band (12-30Hz) for the stimulated data, where no significant difference was observed (Figure 6 A, B).

**Figure 6.**
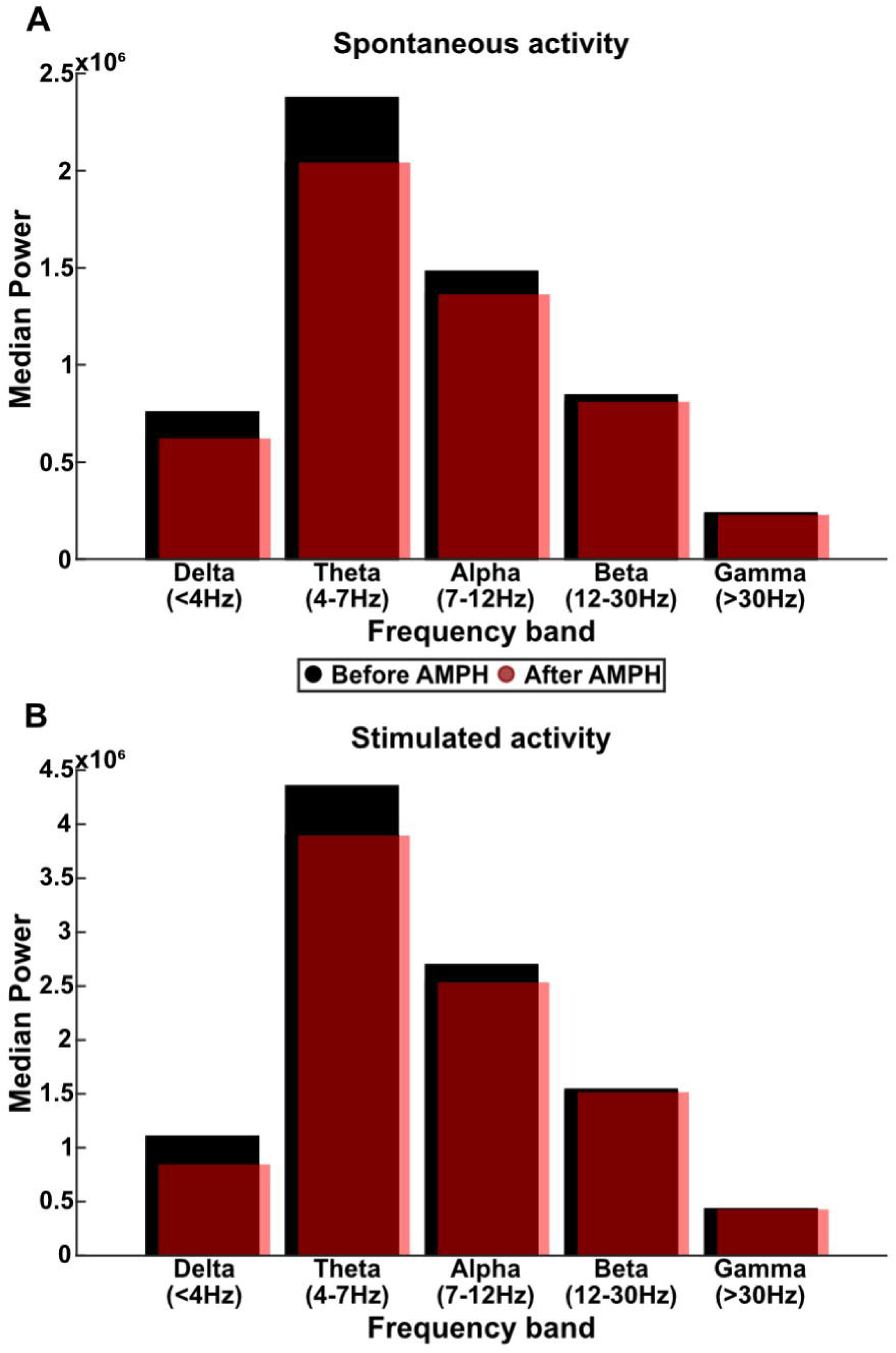
D-amphetamine reduced the power across all frequency bands. **A** The power in each frequency band during all spontaneous activity from one experiment, divided into before (black bars) and after (opaque, red bars) the administration of D-amphetamine. **B** Similar to **A** but for the stimulated activity.

## Discussion

D-amphetamine significantly altered the activity distribution patterns recorded by multi-channel ECoG activity. This was true for both spontaneous activity and for brain activity in the presence of ongoing sensory input. We also found that the difference between the spontaneous and the stimulated activity was decreased by D-amphetamine. As discussed in greater detail below, our analysis indicates that D-amphetamine altered the cortical activity distribution patterns, equivalent to that D-amphetamine pushed the cortical network activity towards new locations in its state space. This is a new angle of how to interpret the effects of D-amphetamine on cortical activity dynamics, extending the more traditional descriptions that it impacts dopamine, serotonin, noradrenaline and acetylcholine transmitter systems or specific aspects of averaged functional connectivity.

### D-amphetamine impacts the preferred state space locations of the network

The impact of D-amphetamine was quantified using a previously published approach[5], where the activity distribution patterns across multiple ECoG recording electrodes are used as an estimation of changes in the global activity distribution patterns across sets of neuron populations. The changes in the neural activity distributions, which occurred for every 1 ms-time step of the recording data, can be regarded as a proxy of the changes in the global state of the network. As previously described, the ‘realm’ of possible activity distributions in the global neuron population will form a high-dimensional state space (see for example [23]). Whereas the unperturbed network activity may normally have preferred and non-preferred locations in that state space[23], here we wanted to quantify if D-amphetamine impacted the preferred state space locations of the brain activity, similar to what has been observed for tactile and visual inputs[5,22] and for auditory inputs[23]. Since we have previously shown that ongoing, weak tactile stimulations can induce systematic changes in state space locations, we quantified the impact of D-amphetamine on both spontaneous activity and on activity recorded during ongoing tactile stimulation. The findings that D-amphetamine perturbed both types of activity (Figure 2) and that the difference in the state space locations between the spontaneous and the stimulated activity was reduced (Figure 3), both effects of which occurred across all dimensions of the activity state space (Figures 4, 5), indicate that D-amphetamine also resulted in a loss of information about real-world inputs in the brain circuitry activity. Altogether, this suggests that one effect of D-amphetamine was a reduced structure in the brain activity, such that the stimulated activity was less separated from the spontaneous activity in each dimension. Less structure in each dimension of the brain activity data would imply a more chaotic temporal evolution of the brain activity, potentially implying a less predictable or controlled behavior which could potentially explain some of the symptoms exhibited by people using amphetamine as a drug[13].

It should be noted that the observation that each dimension of the data, i.e. each feature in the ECoG activity distribution patterns, carried similar weight in explaining the differences induced by D-amphetamine (Figures 4,5) argues against any simple interpretation such as an amplification of the thalamocortical loop activity. If D-amphetamine’s effect would simply have been an impact on the general excitability of the thalamocortical loop, then it would have been expected to result in more prominent effects in one or a few dimensions of the data. Rather, this observation supports that the alterations induced by D-amphetamine impacts the physiological network structure in multiple ways, such that new pathways of activity spread across the network opens up[24], and that this is what caused the network to find new preferred locations in its activity state space.

However, since our analysis method is very sensitive to fine nuanced changes in brain activity, we used comparatively low doses of D-amphetamine. It is conceivable that at higher doses there may exist dose-dependent reductions in the dimensionality of the brain activity. Other studies of the effects of D-amphetamine on EEG recordings detected an effect only after repeated administrations of a higher doses (0.6mg/kg I.P. as opposed to 0.25mg/kg I.V. used here) and these effects may possibly involve additional factors compared to those recorded in the present study[10].

### On the probability of obtaining clustered data with the kNN method

Because the pattern of the recorded multichannel ECoG signal will be constantly changing due to ongoing internal processes in the brain networks, even comparing two separate segments of spontaneous activity under the same condition will have a certain probability to be reported to be different by the PCA-kNN approach we used. This can be seen in Figure 2 C,D, where different half-segments of recorded ECoG data recorded under the same condition (i.e., with or without D-amphetamine) were found to be different by the kNN analysis. This effect would be expected to be time-dependent, i.e., if we had recorded spontaneous activity for 48 hours and just compared the first 24 hours to the second, the kNN accuracy would have been more likely to be closer to chance (50%). Conversely, the accuracy of such a comparison would have increased if we instead compared 10-minute segments and increased even more if we compared just one-minute segments to each other. This is not surprising but merely indicates how sensitive the method is and how complex the brain activity patterns normally are[25–27].

### The effect of D-amphetamine on the distribution of ECoG frequency power

A potential explanation for the differentiation of activity before and after D-amphetamine could be the altered frequency content of our ECoG signal. The ECoG primarily signals local field potentials, which in turn primarily reflects synaptic activity. Field potentials thereby indicate changes in the activity in the underlying neuron population[28]. The broad-band loss of power in all frequencies would suggest that there were fewer or less synchronized activity changes in the neuron population after amphetamine. The only exception was the Beta band for the activity recorded during ongoing tactile stimulation. Previous studies of the effect of amphetamine across EEG frequency bands have reported a reduction in slow wave activity after administration of amphetamine using a lower dose than here (0.15mg/kg, I.V.)[11]. A similar reduction in slow wave activity, together with an increase in alpha band frequency was seen after using a higher dose of D-amphetamine than used in this study (0.6mg/kg, I.P.)[10]. Similarly, another study found that a low dose of D-amphetamine (0.4mg/kg I.P.) causes a desynchronization with general lowering of power in all frequency bands, while a high dose (4mg/kg I.P.) increased power at the 7 to 9.5 Hz range (alpha-1 band), likely due to different receptors being affected at different dosages [29]. This decrease in power in all frequency bands is in agreement with our findings. The lack of significant effect of D-amphetamine on the stimulated activity in the Beta band (Figure 6) could potentially be a sign of our stimulation having an amphetamine-dependent effect on those specific frequencies, though the stimulation frequencies were disjunct from this band.

Previous studies have reported a decrease in slow-wave activity during wakefulness following oral D-amphetamine administration, accompanied by a reduction in both non-REM and REM sleep duration[18]. I.V. administration of D-amphetamine at doses of 0.3-3mg/kg has been shown in rats to induce a dose-dependent arousal from sevoflurane anesthesia, reduce the time to emergence from propofol anesthesia, and accelerate the recovery of consciousness and respiratory drive following fentanyl administration. Its effects during ketamine anesthesia appear to differ, however. One study found that while I.V. D-amphetamine at a dose of 1 mg/kg did reduce the time to emergence after dexmedetomidine administration, a higher dose of 3mg/kg did not significantly reduce the time to emergence from ketamine anesthesia[30–32]. This means that while D-amphetamine have a general arousal effect, this effect seems to be reduced or removed in some way during ketamine anesthesia, potentially due to the fact that ketamine can increase the release of dopamine in the prefrontal cortex[33,34] and thereby interfere with some of the effects that amphetamine is expected to have on this transmitter system.

### Limitations of the field potential approach to analyze brain activity dynamics

Could coarse mass electrode recordings from some perspectives offer advantages compared to multi-neuron recordings? Neuron recordings naturally have a higher resolution, but the ECoG recordings are naturally more globally distributed and more easily conducted. It is not theoretically possible to record from every single neuron in the brain, in fact the tissue-destructive effects of inserting electrodes into the brain tissue limits the single neuron approach to record from extreme subsets of the entire neuron population. If cortical operation is the effect of globally integrated signals, the more distributed signal pickup may provide at least some advantages compared to the details of local signals. With the major disadvantage of course being that the recorded transitions in neuron population activities become extreme under-representations of which specific neurons alter activity in which direction. However, recent work using multichannel ECoG recordings to drive a diversified speech synthesizer indicate that this type of recording can indeed contain a lot of information about the underlying brain processing[35]. Moreover, ECoG is closely related to non-invasive EEG, and the results we obtained here are likely to be translatable to that non-invasive method, which could hence be applied also in humans.

### Concluding remarks

D-amphetamine altered the global patterns of brain activity distributions for both spontaneous and stimulated activity. This change in activity distributions was also associated with a reduction of the difference between spontaneous and stimulated activity. This implies that amphetamine caused a reduction in the organization of the brain’s network activity, in the sense that internally generated brain activity became less separable from the brain activity in the presence of sensory stimuli.

## Methods

### Ethical considerations

Animal experiments were approved in advance through the Local Animal Ethics committee in Malmö/Lund (permit ID M13193-2017 and M20013-2021with addendum V2023/154). All experiments were carried out in accordance with local laws and guidelines. All methods have been carried out in accordance with ARRIVE guidelines.

In this paper we apply the same analysis methodology as in a previous paper[5], but with other tests to understand the impact of D-amphetamine on detecting and decoding sensory input.

### Animals and preparation

Adult Sprague-Dawley rats (male, n=14, 314-611 grams) were initially sedated using isoflurane (3% mixed with air for 60–120 s), and then anesthetized using a ketamine/xylazine mixture (ketamine: xylazine ratio of 15:1, initial ketamine dose of 60mg/kg) that was injected intraperitoneally, before undergoing preparatory surgery. Once catheter was inserted into the femoral vein, anesthesia was maintained using a continuous intravenous infusion of a ketamine/xylazine mixture (ketamine: xylazine ratio of 20:1, initial ketamine dose of 5mg/kg). A second intravenous catheter was inserted in the femoral vein on the other side to allow the administration of D-amphetamine sulfate without interrupting the anesthesia. To ensure a deep enough level of anesthesia we controlled for the absence of withdrawal reflexes in response to noxious pinch to the hind paw. Once the recording of ECoG activity had been established, the occurrence of episodic slow wave sleep activity was used together with the lack of withdrawal reflexes to continue monitoring sedation level.

We made four 5×5 mm craniotomies to access the cortical recording areas, two over the sutura coronaria to access the primary motor and somatosensory areas, and two rostrally to the sutura lambdoidea to access the primary visual and auditory cortex (Figure 1A). To keep the brain surface moist, a cotton and agar pool was made and filled with 37[paraffin oil covering the exposed cortical areas. The dura mater was cut in the rostral part of the exposed areas to allow the escape of cerebrospinal fluid CSF and ensure the dura laid flat against the cortical surface. Cotton drains over the pool edge were used to continuously drain the CSF. Eight silver ball electrodes (diameter 0.25 um) were placed in the eight recording areas (Figure 1A) to record ECoG activity, and two grounding electrodes were placed in the neck muscle. The two electrodes in the primary somatosensory area (S1) were placed in the forelimb area.

### Recordings

A Digitimer NL844 pre-amplifier with low frequency cut off at 0.1Hz and gain x1000 and a NL820 isolator (Neurolog system, Digitimer) with gain x5 amplified the recording signal and fed it into a CED 1401 mk2 hardware, digitizing the voltage data at 1 kHz. The digitized signals were visualized and saved to hard drive using the Spike2 software (Cambridge Electronic Design (CED), Cambridge, UK). Local field potential responses evoked by the skin stimulation verified the placement of S1 electrodes in the forepaw area. No experiment lasted longer than eight hours after the induction of the anesthesia and all animals were euthanized using pentobarbital.

### Stimulation protocol

Two pairs of intracutaneous needles were used for electrical skin stimulation. They were inserted at the base of the second digit of the left forepaw and of the right hind paw, respectively (Figure 1B). The stimulation was delivered as single pulses to one stimulation site at a time (pulse intensity 0.5 mA, pulse duration 0.14 ms). The pulse intensity was set to be above the threshold for activation of tactile afferents while still below the threshold for pain fibers[36,37]. The stimulation protocol alternated periods of spontaneous and stimulated activity, always starting and ending with spontaneous activity. The stimulated periods consisted of repeated single pulse stimulations to a paw at a set frequency of either 0.3, 0.5, 1, 2, 3, 4 or 5 Hz lasting for 5 minutes. Each spontaneous period lasted 2 minutes. Each frequency of stimulation was first delivered to the left forepaw, and then, after a period of spontaneous activity, to the right hind paw, before the frequency was increased for the next period of stimulation. D-amphetamine was administered once the first run of the protocol had been completed. 15 minutes after the administration the same stimulation protocol was repeated (Figure 1C).

### Drug administration

D-amphetamine sulphate solution (Sigma-Aldrich, SE) with a salt weight of 1mg/ml was injected intravenously at a dose of 0.25mg/kg, with a vehicle of 0.3ml of 0.9% saline solution. The level of anesthesia was closely monitored following the injection, to ensure the animal did not wake. If needed, the infusion rate of the anesthesia was increased to keep the animal properly anesthetized.

### Data collection

9 animals were used for the D-amphetamine protocol. The ECoG activity from 7 of 9 of these animals were analyzed. Two experiments were excluded due to difficulties with the recording quality and D-amphetamine was never administered in these cases. The stimulation protocol lasted about two hours with alternating periods of spontaneous and stimulated activity, as described above (Figure 1C), and was repeated twice, once before and once after the administration of D-amphetamine. In addition, in 5 different animals we made control experiments for the same duration, but without administering the D-amphetamine.

### Post processing

The ECoG data was imported to MATLAB (MATLAB Release 2021a, The MathWorks, Inc. Natick, Massachusetts, United States) from Spike2. Artifact removal was done using linear interpolation between the two time-steps just before and after the time of the stimulation impulse. To smooth the raw ECoG data, a Savitzky-Golay filter with a window size of 20 ms was applied (Figure 1D).

### Principal component analysis

To analyze the activity distribution patterns of the Z-scored ECoG data at each time step, principal component analysis (PCA) was used. The principal component (PC) vectors were calculated from the complete ECoG dataset (both before and after D-amphetamine administration) using MATLAB’s inbuilt function “*pca*”.

### kNN analysis was used to quantify data in two types of comparisons

K-nearest neighbor (kNN) was used to compare the distribution of data points in the high-dimensional PC space. The comparison was made on pairs of data sets, such as spontaneous activity being compared with activity under ongoing stimulation. kNN then provided a measure of how often, or with what probability, neighboring data points were of the same category. A kNN output above 50% would indicate a degree of separation between the two sets of data. We made two fundamentally different types of tests. One test compared the spontaneous activity before and after D-amphetamine, and for the same test we also tested if the activity during ongoing tactile stimulation was altered by D-amphetamine (Figure 2). The other type of test was instead using kNN to quantify the difference between spontaneous and stimulated activity before amphetamine and then testing if the kNN result for the difference between spontaneous and stimulated activity was greater or smaller after amphetamine (Figure 3).

Since there were 7 stimulation frequencies from 2 sites across 7 animals, a total of 98 kNN results were obtained for each test. Note that this also applied when we quantified the spontaneous activity, as there were 98 segments of spontaneous activity between the different stimulations.

### kNN analysis

The k-nearest neighbor (kNN) analysis was applied to the PCA coefficients of each data point in the comparison. This was done using MATLABs classification learner toolbox with N=5 nearest neighbors and five-fold cross-validation, in the same manner as in Mellbin et al. 2024[5]. Each time series of PC coefficients was normalized to ensure that that also the higher order PCs could impact the kNN result.

Stimulated activity was defined as the 190 ms time window that followed the preceding stimulation pulse, to avoid overlap with the stimulation pulses for the 5Hz stimulation. As a consequence, at lower stimulation frequencies, the period of stimulated activity contained less data points than the full period of spontaneous activity (see “Stimulation protocol” above). In such cases, we used a shorter, random consecutive range of data points from the spontaneous activity to obtain the same number of data points when comparing spontaneous and stimulated activity. For each comparison, kNN analysis was repeated 100 times, to get a mean decoding accuracy. Together with the five-fold cross-validation this means that each reported decoding accuracy was the result of 500 randomizations of test and training data.

When comparing matching periods of the stimulation protocol before and after D-amphetamine administration, the period of activity was divided into two halves and the kNN was performed 50 times on each half for a total of 100 runs, with the accuracies for all of these runs then being grouped (Figure 2A). We also controlled if the separability of the same type of activity after D-amphetamine was significantly different than the separability of the same type of activity within the same condition (either before D-amphetamine or after D-amphetamine). The control was done on a single period instead of between two different periods to remove any effects from different stimulation frequencies, stimulation placements and preceding periods (Figure 2A). When comparing spontaneous and stimulated activity, a new time series from the spontaneous activity was randomized every 10^th^ iteration of the 100 repetitions, to avoid a specific subset of spontaneous activity skewing the result.

To investigate the significance of the decoding accuracies, each kNN analysis was also repeated on shuffled data. The shuffled data contained the same data points but with shuffled group labels (stimulated or spontaneous activity/pre or post D-amphetamine). Shuffling was repeated 100 times. A full analysis procedure was applied to the shuffled data, with the full set of 100 shuffles.

For significance testing when comparing the kNN activity before and after D-amphetamine to two control groups, Friedmans test was used, and the sign test was used as a post hoc test. We also investigated how the result of the kNN analysis altered when excluding individual PCs one-by-one from the analysis, and by varying the number of PCs included, from one to eight.

### Frequency analysis

To analyze the effect of D-amphetamine on the frequency content of the ECoG data, we applied a continuous wavelet transform on the filtered ECoG data using MatLab function *“cwt”* with default parameters. This was done both for the whole two-hour period before and two-hour period after D-amphetamine administration, as well as for the spontaneous activity (two-minute periods, n=98) and the stimulated activity (five-minute periods, n=98) before and after D-amphetamine. We calculated the power in the following frequency bands: Delta <4Hz, Theta 4-7Hz, Alpha 7-12Hz, Beta 12-30Hz and Gamma >30Hz. Sign test was applied to the paired data to test for any significant differences in the frequency content before and after D-amphetamine administration.

## Data availability

Original ECoG data reported in this paper will be shared by the lead contact upon request. This paper does not report original code.

Any additional information required to reanalyze the data reported in this paper is available from the lead contact upon request.

## Acknowledgments

This work was funded by the Swedish Research Council VR, project no. 2019-01623.

## Author contributions

AM and FB carried out experiments. AM did the formal analysis, wrote the original draft and prepared the figures. All authors were involved in developing the methodology and reviewing the manuscript.

## Additional information

The authors declare no competing interests.

